# Co-Circulation of Multiple Kolmiovirus Lineages Through Vertebrate Evolution

**DOI:** 10.64898/2026.06.05.730526

**Authors:** Lauren Lim, Kate van Brussel, Julien Mélade, Miriam Boucher, Benjamin B. Parrott, Sarah L. Whiteley, Ethan Mandojana, Thomas R. Rainwater, James T. Anderson, Karrie Rose, Edward C. Holmes

**Affiliations:** School of Medical Sciences, the University of Sydney, Sydney, NSW 2006, Australia; Belle W. Baruch Institute of Coastal Ecology and Forest Science, Clemson University, Georgetown, South Carolina, USA; Savannah River Ecology Laboratory, University of Georgia, Aiken, SC, USA; Eugene P. Odum School of Ecology, University of Georgia, Athens, GA, USA; Institute for Applied Ecology, University of Canberra, Bruce, Australia; Department of Molecular Biology, Princeton University, Princeton, NJ, USA; Tom Yawkey Wildlife Center and Belle W. Baruch Institute of Coastal Ecology and Forest Science, Clemson University, PO Box 596, Georgetown, SC 29442, USA; Australian Registry of Wildlife Health, Taronga Conservation Society Australia, Mosman, NSW 2088, Australia

**Author notes:** Prof. Edward C. Holmes **Email:**.

**Keywords:** Deltavirus, *Kolmioviridae*, Evolution, Phylogeny, Co-Divergence

## Abstract

Although once only characterized by human hepatitis deltavirus (HDV), membership of the family *Kolmioviridae* has dramatically expanded in recent years. Despite this transformation in our understanding of the host range of kolmioviruses, the evolutionary history of this enigmatic group of RNA viruses is unclear. Kolmioviruses are characterized as small (∼1.7kb) satellite viruses that encode a single ∼200 amino acid delta antigen (DAg) and require unrelated helper viruses for replication. Here, we describe eight novel kolmioviruses from metatranscriptomic studies of the American alligator (*Alligator missippiensis*), red kangaroo (*Osphranter rufus*), and central bearded dragon (*Pogona vitticeps*), as well as avian kolmioviruses mined from the Sequence Read Archive (SRA). Although the novel kolmioviruses were often found in samples co-infected by other viruses, there was no evidence for the presence of hepatitis B virus as seen in HDV. By employing a range of sequence data sets, alignment methods, alignment trimming methods, and substitution models, we provide an evolutionary history of the *Kolmioviridae* that maximizes the extent of virus-host co-divergence and refines estimates of their evolutionary timescale. Although DAg amino acid sequences are more conserved than nucleotide sequences and hence might be expected to result in more accurate phylogenetic trees, we show that full genome nucleotide sequences likely provide the best representation of kolmiovirus evolution. More broadly, our results reveal that irrespective of the data set used, multiple distinct kolmiovirus lineages have co-circulated throughout vertebrate evolution over timescales spanning hundreds of millions of years, with the association between HDV and HBV appearing only recently.

**Significance Statement:** Kolmioviruses are satellite RNA viruses, with hepatitis deltavirus (HDV) associated with human disease following co-infection with hepatitis B virus (HBV) the best characterized. Although a growing number of animal kolmioviruses have been identified in metagenomic studies and associated with a range of helper viruses, the evolutionary origins and history of this important and unique group of viruses is unknown. By identifying novel kolmioviruses in a range vertebrate hosts, including American alligators, we show that highly diverse lineages of kolmioviruses have co-circulated for the duration of vertebrate evolution with clear evidence of virus-host co-divergence, and are associated with a variety of potential helper viruses. Despite this antiquity, we present evidence that the HDV-HBV association only recently evolved in human populations.

## Introduction

The *Kolmioviridae* are a family of unusual circular negative-sense RNA viruses that use a positive-sense RNA antigenome as a replication intermediate (1). Kolmioviruses are characterized by small genomes (∼1700 nucleotides in length), the presence of a single protein – the delta antigen (DAg) – as well as a non-coding catalytic RNA molecule that functions as a ribozyme in virus replication, and are unrelated to other RNA viruses such that they are classified within their own realm (Ribozyviria). To produce infectious particles, kolmioviruses are thought to rely on a coinfecting enveloped virus as a helper (1). As a case in point, the best characterized, and until recently the only known kolmiovirus – human hepatitis deltavirus (HDV; genus *Deltavirus*) – utilizes the surface antigen of hepatitis B virus (HBV) to produce virus particles, such that HDV is considered a satellite virus of HBV (2, 3). Considerable attention has been directed towards understanding the consequences of HDV and HBV co-infection as it is associated with a worse clinical outcome, most notably fulminant hepatitis (4).

Until 2018 human HDV was the only known deltavirus. As a consequence, the evolutionary origin of this virus was largely mysterious, including suggestions that it was related to plant viroids that similarly possess ribozymes (5) or represented an escaped human intron (6). The first indication that viruses related to HDV were present in other animals was their discovery in birds (7) and snakes (8), with the latter successfully isolating the virus (denoted SDeV). Since these initial discoveries, the large-scale generation and analysis of metagenomic data has led to the identification of “delta-like agents” in other animal taxa, including placental mammals (such as bats and rodents), marsupials, birds, amphibians, fish, and invertebrates, as well as divergent relatives of these viruses in environmental samples (9–12). In 2024, these “delta-like agents” were classified within the newly established *Kolmioviridae* family by the International Committee on Taxonomy of Viruses (ICTV) (13).

Simultaneous with an expansion of host range, metagenomic data have broadened both the range of kolmiovirus tissue tropism beyond the strict hepatotropism observed in HDV and the range of helper viruses beyond HBV. Although the new mammalian kolmioviruses contain a small delta antigen (SDAg), they lack the large delta antigen (LDAg) present in HDV (14). The LDAg is characterized by a 19-20 amino acid extension at its C terminal that contains a prenylation motif (CXXQ) thought to play a crucial role in its interaction with both the ribonucleoprotein and hepatitis B surface antigen (HBsAg). The absence of both LDAg and HBV, along with the presence of other candidate helper viruses, suggests that most kolmioviruses do not rely on co-infection with HBV or other hepadnaviruses to replicate (15). In addition, metagenomic sequencing has highlighted potential helper viruses such as bornaviruses for some avian kolmioviruses (12), and experimental studies with SDeV have shown that reptarenaviruses and hartmaniviruses (*Arenaviridae*) can successfully act as helpers in the production of infectious SDeV particles (16), while influenza viruses, hepaciviruses, herpesviruses, poxviruses, and retroviruses have also been proposed as helpers (1). Similarly, recent studies of rodent kolmioviruses indicate that they are able to utilize a range of helper viruses via co-packaging, including rhabdoviruses, herpesvirus and arenaviruses (17). It has also been suggested that some kolmioviruses, particularly fat-tailed dunnart deltavirus, may not need helpers at all (18).

As well as exogenous kolmioviruses, endogenous relatives of these viruses have been identified in termite genomes, with molecular clock dating suggesting that they have existed for approximately 141 million years (19). The same study also estimated that the evolutionary timescale of the exogenous vertebrate kolmioviruses was less than 240 million years, comprising a phylogenetic history characterized by frequent cross-species transmission (i.e. host jumping). Indeed, most studies of kolmiovirus evolution have identified relatively frequent cross-species transmission in comparison to virus-host co-divergence.

Although there has been a revolution in our understanding of the diversity and biology of kolmioviruses, the fundamental patterns and processes of kolmiovirus evolution remain poorly understood. Despite a growing list of kolmioviruses, the evolutionary history of this family has proven challenging to resolve, and the apparent lack of virus-host co-divergence has made the time-scale of their evolutionary history difficult to determine. At the time of writing, the *Kolmioviridae* comprises 11 genera, within which there is usually only a single species (with the genus *Deltavirus* that contains eight species [also referred to as genotypes] of human HDVs the most notable exception) (13). That there are so many genera with so few component species points to an extensive unsampled diversity of kolmioviruses. Although the current host species distribution of exogenous and endogenous kolmioviruses suggests that this family is ancient, there is no consensus on the timescale of their evolutionary history. However, as these viruses are highly divergent in sequence and found in vertebrate species that themselves diverged hundreds of millions of years ago, it would be surprising if there were no instances of virus-host co-divergence over this long evolutionary timespan.

Very high levels of sequence divergence among viruses, which results in substantial site saturation (i.e., an excessive number of nucleotide and/or amino acid substitutions per site) in evolutionary comparisons, compromises phylogenetic analysis. This is particularly acute in the case of the kolmioviruses, in which very small genomes necessarily mean that there are few available nucleotide or amino acid sites for phylogenetic analysis. In addition, the phylogenies generated to date suggest substantial rate variation in kolmiovirus evolution, manifest in tree branches of greatly varying length (1). This may compromise the accuracy of both phylogenetic analyses and molecular clock dating, and is indicative of a substantial unsampled genetic diversity.

To help resolve the evolutionary history of the *Kolmioviridae*, we identified novel kolmioviruses from ongoing metatranscriptomic sequencing projects of diverse animal species and mined them from the Sequence Read Archive (SRA). With these newly acquired kolmioviruses we performed phylogenetic analyses using a variety of data sets and substitution models to better assess the extent of co-divergence across the kolmiovirus phylogeny and to more accurately reconstruct the pattern and timescale of virus evolution.

## Results

### Identification of kolmioviruses in American alligator, central bearded dragon, and red kangaroo

As part of ongoing studies in virus ecology and evolution we have generated RNA sequencing libraries from diverse animal taxa. Herein, we screened these data for kolmioviruses. Accordingly, kolmioviruses were identified in six sequencing libraries: three from the American alligator (*Alligator mississippiensis*), two from the central bearded dragon (*Pogona vitticeps*), and one from the red kangaroo (*Osphranter rufus*). Metatranscriptomic data sets ranged from 71,834,568 to 140,540,704 paired-end reads and 21-50 Gb of sequence data per library.

Metagenomic screening using Kraken2 revealed no unexpected species or human contamination in any of the libraries, with each containing sequence reads reflecting the host in question as well as those likely associated with the environment or host diet. Host reads ranged between 88.33-99.95% within the red kangaroo and central bearded dragon tissue libraries, while the American alligator pools largely contained *A. mississipiensis* reads but with greater variation (45.66-91.85% host reads). This wider range likely resulted from the addition of oral and cloacal swabs which introduced more environmental bacterial reads into the sequencing libraries. In addition, the American alligator and central bearded dragon libraries from which the novel kolmioviruses were identified contained a range of other vertebrate-associated viruses. Specifically, American alligator libraries contained viruses from the orders *Picornavirales*, *Nodavirales* and family *Picobirnaviridae*, while the central bearded dragon libraries had representatives from the order *Picornavirales* and family *Flaviviridae*. Although these could conceivably act as helpers, confirmation requires additional experimental study beyond the scope of this study. No other vertebrate associated viruses were observed in the red kangaroo library.

American alligator kolmiovirus 1 was found in two libraries from alligators sampled in different USA states (South Carolina and Alabama). Phylogenetic analysis revealed that this virus grouped with a lineage associated with bird kolmioviruses, as expected under virus-host co-divergence (Fig. 1). However, the third alligator kolmiovirus sequence, denoted American alligator kolmiovirus 2 (sampled from animals in South Carolina), occupied a different and divergent phylogenetic position, grouping with the viruses found in some amphibians and birds (Chinese fire belly newt, Mallard duck) or ray-finned fish depending on the data set used (Fig. 1; described in more detail below). Given the limited (31%) amino acid sequence similarity between American alligator kolmioviruses 1 and 2, it is likely that they represent two novel kolmiovirus genera. Hence, multiple distinct kolmioviruses can circulate within the same alligator species in the same US state. In contrast, the single central bearded dragon kolmiovirus fell as a sister group to a clade of kolmioviruses found in various bird species, while the red kangaroo kolmiovirus grouped closely with two other marsupials – the Tasmanian devil and the fat-tailed dunnart – forming a marsupial-specific clade.

**Figure 1.**
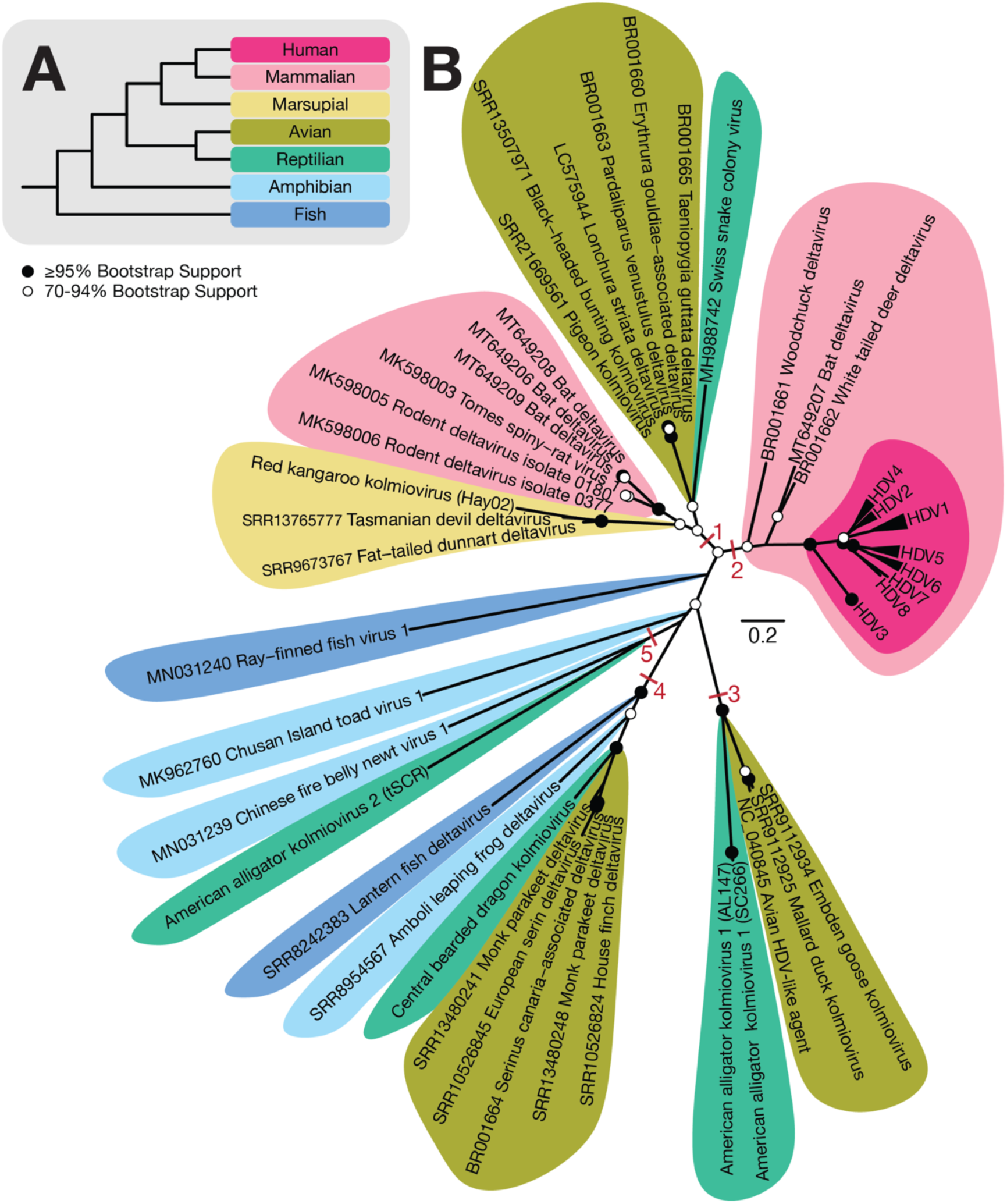
Multiple co-circulating lineages of kolmioviruses. (A) Phylogeny of major vertebrate host taxa color key: human (hot pink), placental mammals (light pink), marsupials (yellow), birds (army green), reptilians (green), amphibians (light blue), fish (dark blue). (B) Unrooted maximum likelihood phylogenetic tree of vertebrate host-associated *Kolmioviridae* based on full nucleotide genome sequence Clustal Omega alignment, trimAl and ModelFinder (TPM2+I+R3) scaled according to number of nucleotide substitutions per site. Nodes 1-5 marked and labelled in red denote lineage clusters compatible with virus-host co-divergence. Nodal support was estimated using 1000 bootstrap replicates, in which nodes ≥95% bootstrap support are annotated with black circles (●) and those 70-94% bootstrap support with white circles (○).

### Identification of avian kolmioviruses in the SRA

Novel avian kolmioviruses were identified in the Mallard duck (SRA library SRR9112925), Embden goose (SRR9112934), Black-headed bunting (SRR13507971), and Pigeon (SRR21669561). Kolimioviruses found in Anseriformes (waterfowl) formed a distinct clade exhibiting strong genome sequence similarity to Avian HDV-like agent (NC_040845) found in a dabbling duck: Mallard duck kolmiovirus exhibited 88% sequence similarity, Embden goose kolmiovirus 87% sequence similarity, while the Mallard duck and Embden goose kolmioviruses shared 96% sequence similarity. Within the avian clade of kolmioviruses, comprising black-headed bunting kolmiovirus, pigeon kolmiovirus, Taeniopygia guttata deltavirus (BR001665), Pardaliparus venustulus deltavirus (BR001663), Lonchura striata deltavirus (LC575944) and Erythrura gouldiae-associated deltavirus (BR001660), full genome sequence similarity ranged between 94-97%. In addition to the novel kolmioviruses, RNA viruses from the order *Nidovirales* and *Flaviviridae* were found in the goose and pigeon libraries, respectively. No candidate vertebrate-infecting viruses were identified within the Mallard duck and Black-headed bunting libraries.

### Revealing the evolutionary history of vertebrate kolmioviruses

Given the diversity of kolmioviruses newly identified here, comprising a number of phylogenetically distinct co-circulating lineages (e.g., American alligators) and indicative of complex virus-host associations, we attempted to better understand the evolutionary history of the vertebrate kolmioviruses as a whole. As the endogenous kolmioviruses found in termites are likely characterized by markedly different evolutionary dynamics than those of exogenous kolmioviruses and comprise a distinct lineage, they were not considered here.

Most studies of kolmiovirus evolution have focused on amino acid phylogenies based on the DAg, particularly as it is the only protein available for analysis and amino acid sequences should be more conserved over evolutionary time than nucleotide sequences. However, these phylogenies are also characterized by highly divergent sequences and both long and variable branch lengths indicative of extensive rate variation, and it is likely that kolmioviruses have experienced variable selection pressures as they are present in diverse hosts. This precluded direct molecular clock dating in the current study. In addition, a major limitation with the use of the DAg in phylogenetic analysis is that the sequence alignment is <200 amino acid residues in length, such that topological resolution intrinsically involves a small number of characters.

To assess the phylogenetic utility of different data sets and substitution models we assumed that the maximum likelihood phylogenetic tree with the greatest number of virus-host co-divergence events was likely the best representation of evolutionary history. In particular, virus-host co-divergence can be reasonably expected to occur if kolmioviruses are present in such diverse classes of vertebrates and likely over extended evolutionary time-scales. Hence, the combination of sequence data set (i.e., amino acid, nucleotide, DAg, full genome) that resulted in the greatest number of co-divergence events (i.e., nodes supporting co-divergence) was considered optimal.

In total, 12 combinations of data set (full genome, DAg amino and DAg nucleotide), alignment method (Clustal Omega, MAFFT and MUSCLE) and substitution model (best-fit or user-defined) were compared in phylogenetic analyses, with the results presented in the first 12 rows in Table 1. To assess the possible impact of outgroup rooting, phylogenetic trees of the full genome, DAg amino and DAg nucleotide sequences with the addition of a single termite outgroup were also estimated (last three rows in Table 1).

**Table 1.**
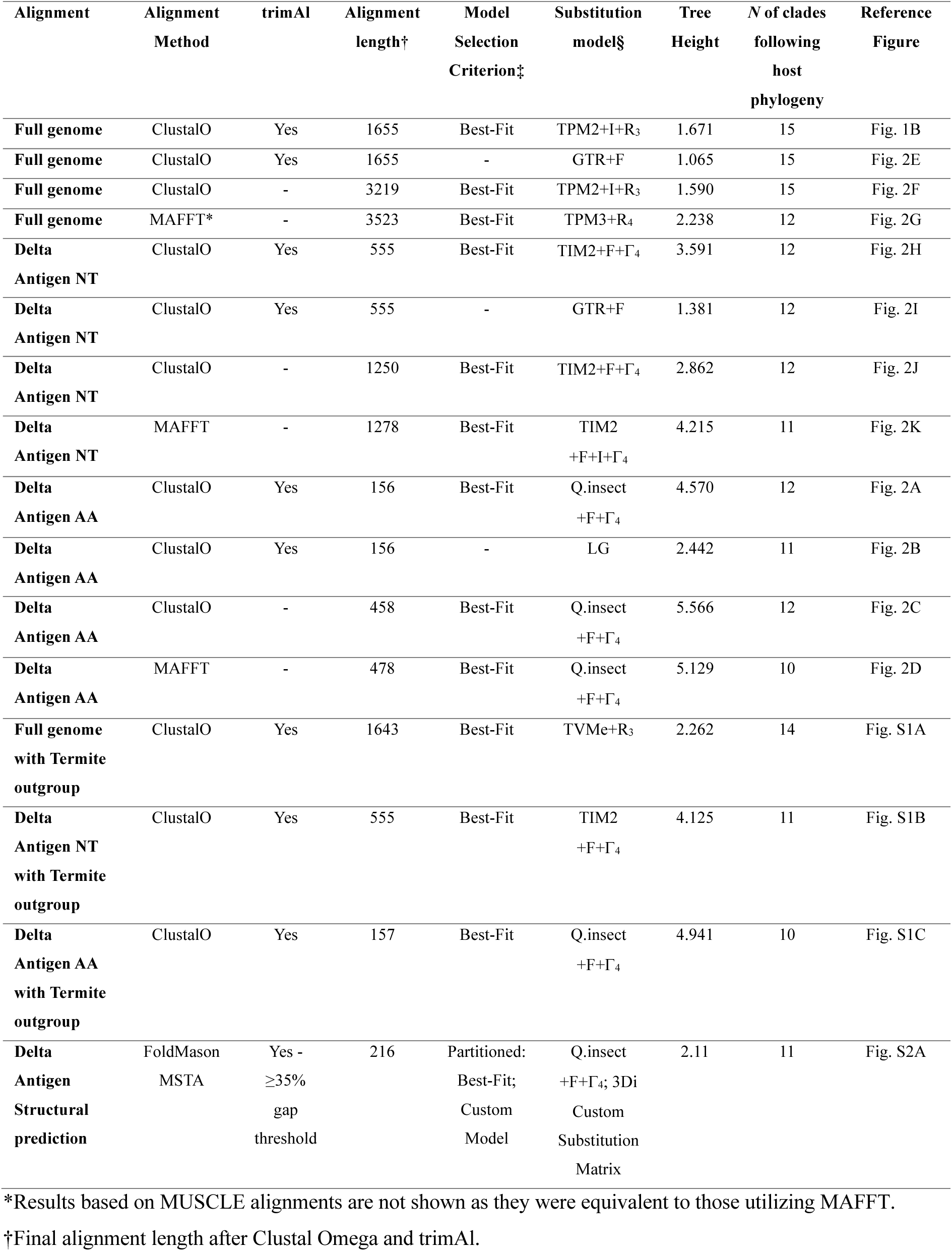
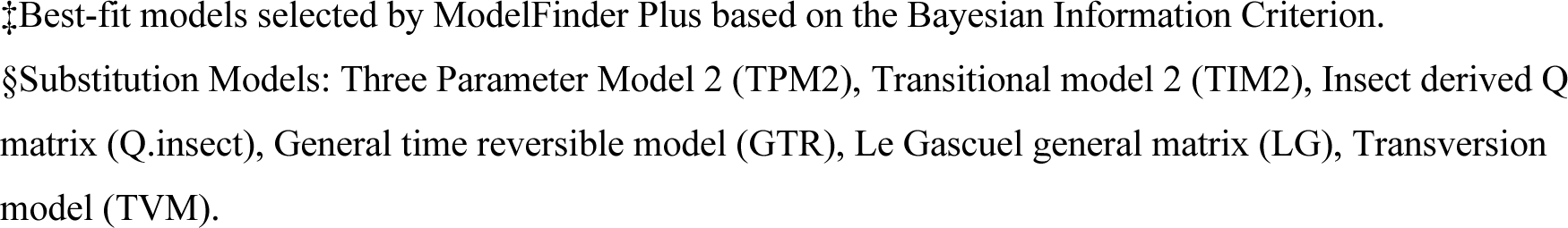
Summary of kolmiovirus data sets and phylogenetic analyses.

| Alignment | Alignment Method | trimAl | Alignment length<sup>†</sup> | Model Selection Criterion<sup>‡</sup> | Substitution model§ | Tree Height | N of clades following host phylogeny | Reference Figure |
| --- | --- | --- | --- | --- | --- | --- | --- | --- |
| Full genome | ClustalO | Yes | 1655 | Best-Fit | TPM2+I+R<sub>3</sub> | 1.671 | 15 | Fig. 1B |
| Full genome | ClustalO | Yes | 1655 | - | GTR+F | 1.065 | 15 | Fig. 2E |
| Full genome | ClustalO | - | 3219 | Best-Fit | TPM2+I+R<sub>3</sub> | 1.590 | 15 | Fig. 2F |
| Full genome | MAFFT* | - | 3523 | Best-Fit | TPM3+R<sub>4</sub> | 2.238 | 12 | Fig. 2G |
| Delta Antigen NT | ClustalO | Yes | 555 | Best-Fit | TIM2+F+Γ<sub>4</sub> | 3.591 | 12 | Fig. 2H |
| Delta Antigen NT | ClustalO | Yes | 555 | - | GTR+F | 1.381 | 12 | Fig. 2I |
| Delta Antigen NT | ClustalO | - | 1250 | Best-Fit | TIM2+F+Γ<sub>4</sub> | 2.862 | 12 | Fig. 2J |
| Delta Antigen NT | MAFFT | - | 1278 | Best-Fit | TIM2<br>+F+I+Γ<sub>4</sub> | 4.215 | 11 | Fig. 2K |
| Delta Antigen AA | ClustalO | Yes | 156 | Best-Fit | Q.insect<br>+F+Γ<sub>4</sub> | 4.570 | 12 | Fig. 2A |
| Delta Antigen AA | ClustalO | Yes | 156 | - | LG | 2.442 | 11 | Fig. 2B |
| Delta Antigen AA | ClustalO | - | 458 | Best-Fit | Q.insect<br>+F+Γ<sub>4</sub> | 5.566 | 12 | Fig. 2C |
| Delta Antigen AA | MAFFT | - | 478 | Best-Fit | Q.insect<br>+F+Γ<sub>4</sub> | 5.129 | 10 | Fig. 2D |
| Full genome with Termite outgroup | ClustalO | Yes | 1643 | Best-Fit | TVMe+R<sub>3</sub> | 2.262 | 14 | Fig. S1A |
| Delta Antigen NT with Termite outgroup | ClustalO | Yes | 555 | Best-Fit | TIM2<br>+F+Γ<sub>4</sub> | 4.125 | 11 | Fig. S1B |
| Delta Antigen AA with Termite outgroup | ClustalO | Yes | 157 | Best-Fit | Q.insect<br>+F+Γ<sub>4</sub> | 4.941 | 10 | Fig. S1C |
| Delta Antigen Structural prediction | FoldMason<br>MSTA | Yes -<br>≥35%<br>gap<br>threshold | 216 | Partitioned:<br>Best-Fit;<br>Custom<br>Model | Q.insect<br>+F+Γ<sub>4</sub>; 3Di<br>Custom<br>Substitution<br>Matrix | 2.11 | 11 | Fig. S2A |

\*Results based on MUSCLE alignments are not shown as they were equivalent to those utilizing MAFFT.
<sup>†</sup>Final alignment length after Clustal Omega and trimAl.
‡Best-fit models selected by ModelFinder Plus based on the Bayesian Information Criterion. §Substitution Models: Three Parameter Model 2 (TPM2), Transitional model 2 (TIM2), Insect derived Q matrix (Q.insect), General time reversible model (GTR), Le Gascuel general matrix (LG), Transversion model (TVM).

Notably, in contrast to expectations based on the impact of multiple substitution, our analysis revealed that nucleotide rather than amino acid phylogenies consistently resulted in a greater number of co-divergence events and hence arguably the best representation of kolmiovirus evolution (Table 1; Fig. 1 and Fig. 2). In particular, the phylogenies estimated from nucleotide sequences of the complete viral genome gave the highest number (n =15) of virus-host co-divergence events, particularly those aligned using Clustal Omega and either untrimmed (alignment length of 3219 nt) or trimmed to remove ambiguously aligned regions (1655 nt), and utilizing either the best-fit models of nucleotide substitution (TPM2+R4 and TPM2+I+R3) or simpler models (GTR+F). These co-divergence events can be grouped into five main nodes (labelled 1-5 on Fig. 1): node 1 – placental mammals, marsupials, birds and reptiles; node 2 – human HDV genotypes/species and other placental mammals (although the grouping among the mammalian species does not itself match the host phylogeny); node 3 – American alligators and birds; node 4 – birds, reptiles, amphibians and ray-finned fish (although this cluster does not yet include mammals as expected under strict co-divergence); and node 5 – American alligators and amphibians. More broadly, this phylogeny indicates that most vertebrate groups – mammals, birds, reptiles, amphibians and fish – harbor multiple and phylogenetically distinct kolmioviruses.

**Figure 2.**
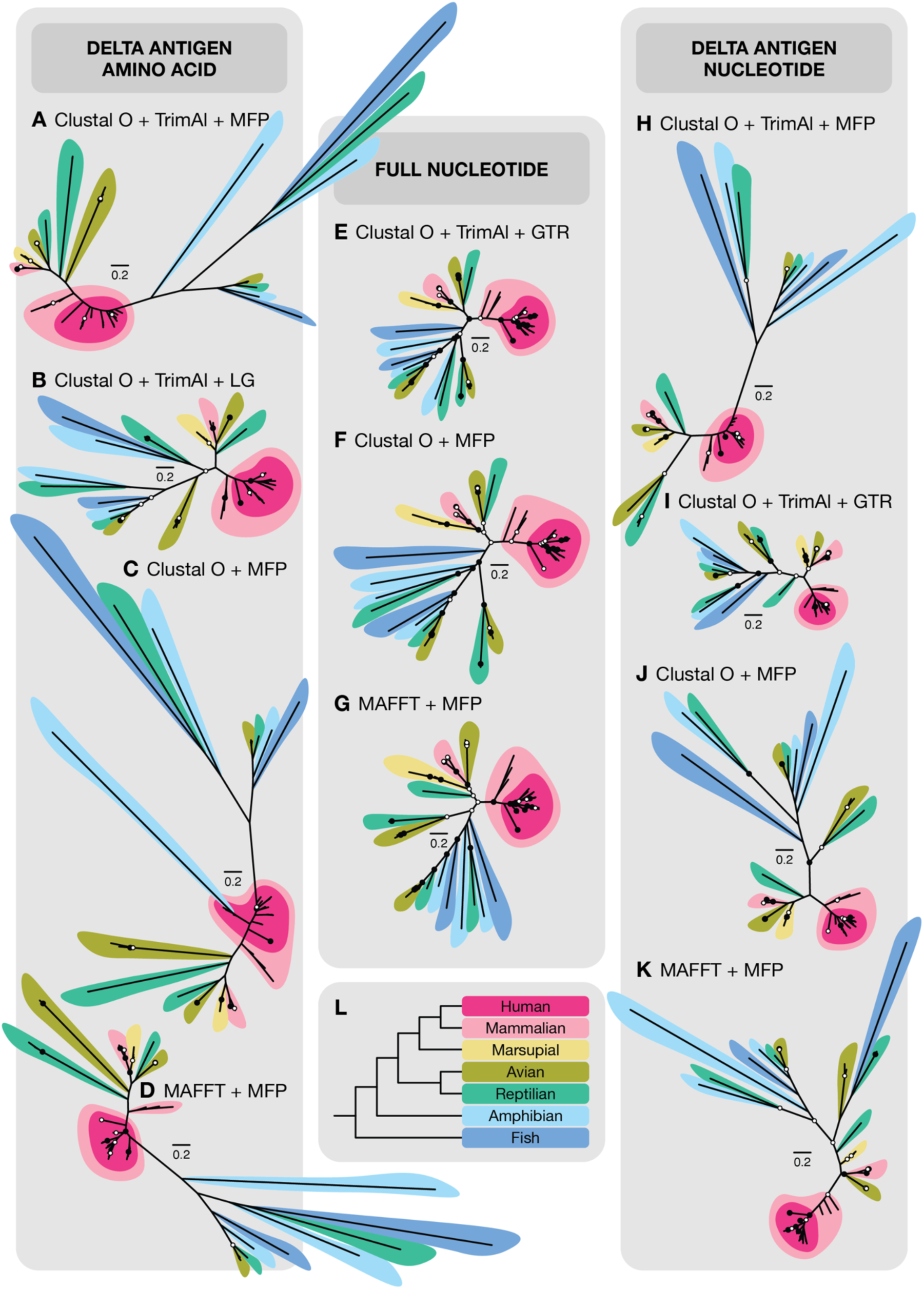
Variation in vertebrate kolmiovirus phylogenies. (A-K) Unrooted maximum likelihood phylogenetic trees of vertebrate host-associated *Kolmioviridae*. Branch lengths are scaled to a common substitution rate such that all panels share a 0.2 substitutions per site scale bar. A-D phylogenies estimated using DAg amino acid alignments, E-G phylogenies using full nucleotide genome sequence alignments, and H-K phylogenies estimated using DAg nucleotide alignments (see Table 1 for details). Nodal support was estimated using 1000 bootstrap replicates; nodes ≥95% bootstrap support are shown with black circles (●) and those with 70-94% bootstrap support with white circles (○). (L) Phylogeny of major vertebrate host taxa color key: human (hot pink), placental mammals (light pink), marsupials (yellow), birds (army green), reptilians (green), amphibians (light blue), fish (dark blue).

Phylogenetic trees based on nucleotide sequences of the DAg resulted in fewer (11–12) co-divergence events than the whole genome phylogeny independent of the method of sequence alignment, whether the alignments were trimmed or untrimmed (which resulted in a >50% reduction in alignment length), or the nucleotide substitution model utilized (Table 1, Fig. 2). In particular, compared to the full genome tree, the phylogeny based on the best-fit model for nucleotide sequences of the DAg commonly lacked support for node 1 that grouped placental mammals, marsupials, birds and reptiles, and in one case (Fig. 2H) even the HDV sequences did not form a monophyletic group.

Although phylogenetic trees based on amino acid sequence alignments of the DAg have formed the basis of most studies on kolmiovirus evolution and classification, they consistently resulted in weaker evidence for virus-host co-divergence and the presence of viral lineages characterized by extremely long branch lengths or unrealistic groupings (Table 1, Fig. 2) compared with nucleotide-based analyses. In particular, the amino acid phylogenies utilizing the best-fit substitution models and with sequences aligned using Clustal Omega and trimmed or untrimmed resulted in 12 instances of co-divergence. In three of the four cases (Fig. 2A, 2C, 2D) the human HDV sequences did not form a monophyletic group (in one case grouping with an amphibian sequence), and the amphibian, fish and reptilian lineages were characterized by markedly longer branch lengths.

To explore why the nucleotide trees are seemingly more realistic than the amino acid trees we estimated the total number of changes per site across each phylogeny, manifest in the total tree height (Table 1). Notably, the tree heights for all the data sets were indicative of large numbers of multiple substitution per site (i.e. >1.0), and those for the DAg amino acid phylogenies where consistently higher than those from the nucleotide phylogenies such that they are revealing more substitutions per site. For example, while the tree height for the best-fit full genome nucleotide phylogeny (Fig. 1) was 1.671, the equivalent for the best-fit amino acid model on the DAg was 2.5 times greater at 4.215 (and that of the untrimmed amino acid alignment attained a tree height of 5.566). As expected, the best-fit, and most complex, models of nucleotide or amino acid substitution resulted in greater tree heights than simpler models, although it was notable that the phylogenetic trees based on MAFFT alignments resulted in greater tree heights and fewer co-divergence events than those generated using Clustal Omega. In sum, while greater tree heights meant that more substitution events are being captured, the very large heights, as well as their substantial variance, indicates that evolutionary distances are routinely and greatly underestimated across the kolmiovirus phylogeny, which will be particularly acute for the longest branches.

Finally, we assessed whether the addition of an outgroup invertebrate (termite) sequence to root the tree improved phylogenetic accuracy (Table 1). Although the whole genome phylogeny including the termite outgroup resulted in 14 co-divergence events, in all cases the termite sequence did not form a longer branch as might be expected from an outgroup lineage such that it did not provide any additional phylogenetic resolution (Fig. S1).

### Phylogenetic analysis based on DAg protein structures

Phylogenetic inference based on the predicted structures of the DAg protein also resulted in a phylogenetic tree with fewer co-divergence events (11) than seen at the full genome level, and with anomalous groupings (Fig. S2). Hence, on these data the AlphaFold3 DAg models lack sufficient structural reliability or confidence for robust phylogenetic inference, in part because of the very short sequence lengths. We also note that the complete structure of the DAg has not yet been resolved and the DAg is regarded as an “intrinsically disordered protein” (IDP) (20, 21). While structural comparisons failed to unveil deeper evolutionary signals within kolmiovirus phylogeny, predictions enabled the identification of structural homology between the well characterized HDV1 and novel kolmioviruses. In particular, there was conservation of the coiled-coil assembly domain and helix-loop-helix (HLH) across the novel kolmioviruses, supported by high structural prediction confidence and structural similarity (Fig. 3). Further disorder prediction indicated that regions of low pLDDT correspond to greater predicted intrinsic disorder (Fig. S3). Considerable variation in the global predicted structure of the LDAg across the *Kolmioviridae* was also observed and likely influenced by these connecting intrinsically disordered regions (IDRs) of low pLDDT.

**Figure 3.**
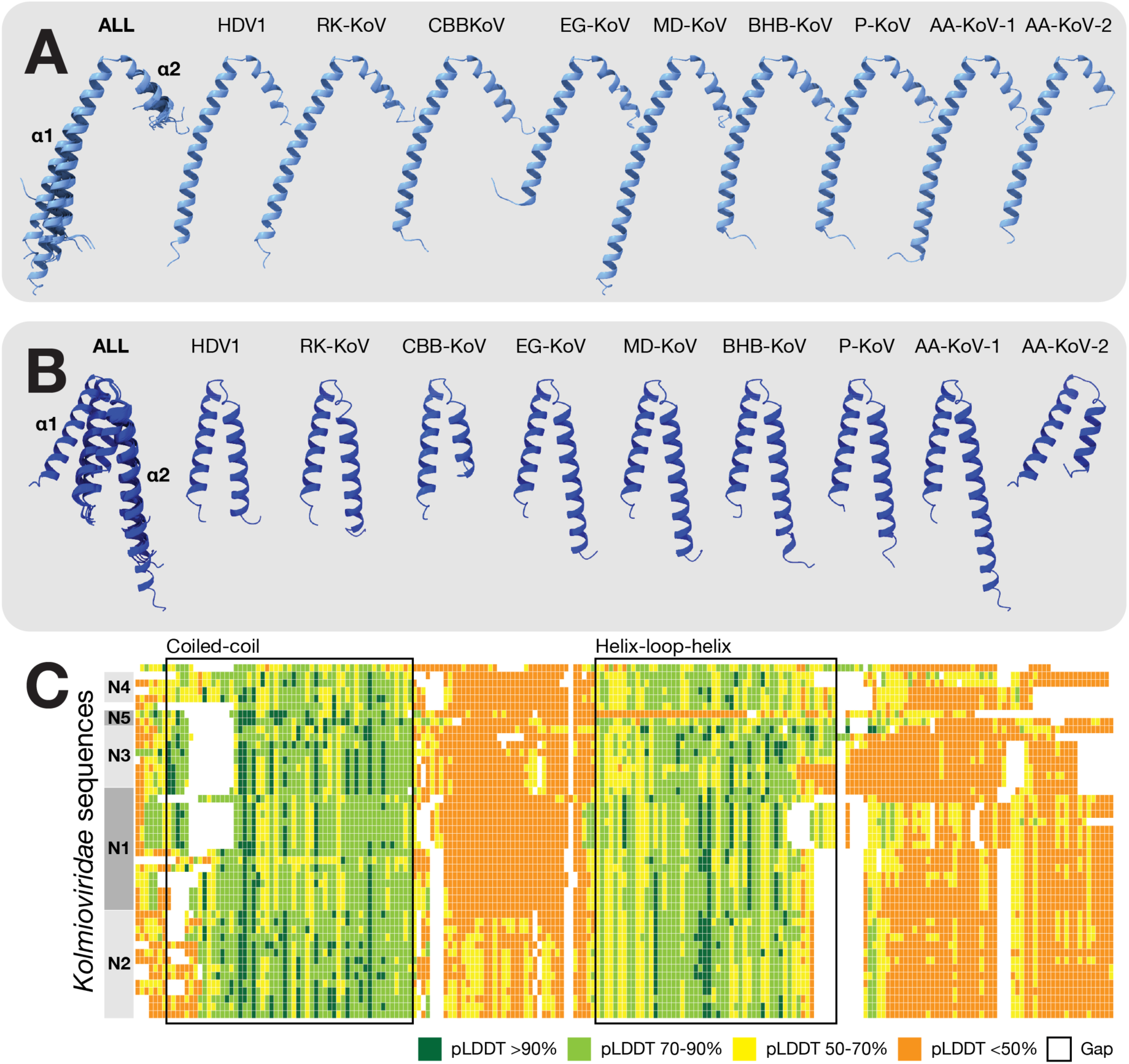
Conserved structural homology in regions of high structural prediction confidence measured by pLDDT in coiled-coil assembly domain and helix-loop-helix domain in novel kolmiovirus DAg. All protein structures are displayed from N to C terminus using ChimeraX. (**A**) Coiled-coil assembly domain in light blue. (**B**) Helix-loop-helix domain dark blue. Labels (left to right): superimposed region (ALL), human deltavirus 1 (HDV1), red kangaroo kolmiovirus (Red-K-KoV), central bearded dragon kolmiovirus (CCB-KoV), Embden goose kolmiovirus (EG-KoV), Mallard duck kolmiovirus (MD-KoV), Black-headed bunting (BHB-KoV), Pigeon kolmiovirus (P-KoV), American alligator kolmiovirus 1 (AA-KoV-1), American alligator kolmiovirus 2 (AA-KoV-2). (**C**) MSTA alignment displayed in a heat map displaying pLDDT, ordered by nodes 1-5 in Figure 1. pLDDT values >90% (dark green), 70-90% (green), 50-70% (yellow), <50% (orange). Approximate region of coiled-coil and helix-loop-helix in alignments outlined.

## Discussion

The increase in our understanding of the phylogenetic diversity of kolmioviruses, coupled with studies demonstrating that the use of HBV as a helper virus is perhaps unique to human HDV, is arguably one of the most revealing stories in the recent history of virus evolution. Herein, we describe four novel kolmiovirus sequences – two from American alligators, and one each from the central bearded dragon and the red kangaroo – and use these sequences to help reveal the evolutionary history of the *Kolmioviridae*. SRA screening also resulted in the identification of four novel avian kolmioviruses, in turn expanding our knowledge of the host range of kolmioviruses in birds. As in other studies of animal kolmioviruses, although additional viruses to kolmioviruses were identified in each of our sequencing libraries, hepadnaviruses were not detected thereby excluding their possible role as helpers. Whether these additional viruses act as helpers requires additional experimental validation.

Despite the increase in the known diversity of kolmioviruses, resolving their phylogenetic relationships and evolutionary timescale has proven more challenging, with current kolmiovirus phylogenies characterized by tree topologies that do not match the evolutionary history of their hosts and that display markedly variable branch lengths. Indeed, our phylogenetic analysis revealed substantial topological uncertainty, with different data sets (i.e., amino acid or nucleotide), gene regions (DAg or complete genome), and substitution models resulting in markedly different phylogenetic trees sometimes possessing highly divergent lineages.

Accurately reconstructing the evolutionary history of the kolmioviruses is evidently complicated by the very high levels of genetic divergence, such that most individual nucleotide and amino acid sites will have experienced (perhaps extensive) multiple substitutions per site. Indeed, the combination of long branches and short sequences is arguably the worst-case scenario for accurately estimating phylogenetic trees. It is therefore notable in this context that phylogenies based on nucleotide sequences, particularly full genomes, provided the strongest evidence for virus-host co-divergence (Fig. 1). Hence, despite the adverse impact of multiple substitution, which will be greater on nucleotide than amino acid sites, it is possible that the greater number of informative sites in nucleotide sequences, particularly at the scale of full genomes, offsets the impact of multiple substitution and provides more phylogenetic resolution as it is less subject to model over-parameterization. In contrast, phylogenetic analyses based on amino acid sequences of the DAg are compromised by a combination of very short sequence alignments, likely model over-parameterization and perhaps poor model-fit, resulting in inaccurate description of kolmiovirus evolution. Indeed, it is striking that the best-fit substitution models for DAg amino acid sequences frequently included a substitution matrix based on insect sequences even though no insects were included in these analyses, and in most cases the human HDV sequences did not form a monophyletic group. In addition, it was notable that phylogenetic analysis incorporating a termite outgroup sequence did not provide any additional resolution and the termite sequence did not fall on a long brancher as might be expected of a true outgroup sequence. This may again reflect the occurrence of widespread site saturation across the kolmiovirus phylogeny.

A key assumption in our study was that the most accurate representation of the evolutionary history of the *Kolmioviridae* was based on a phylogenetic tree that maximized the extent of virus- host co-divergence. Although the relative frequencies of co-divergence versus cross-species transmission vary substantially among viruses (22), co-divergence commonly occurs among major animal classes with cross-species transmission occurring more frequently among closely related taxa that provide similar cellular environments (23). In none of our analyses was there a complete topological match between the virus and host phylogenies, and cross-species transmission has clearly been a common occurrence in kolmiovirus evolution. However, some evidence for co-divergence was apparent in all of our phylogenetic analyses, reflecting phylogenetic patterns that are difficult to explain under cross-species transmission as a whole, and phylogenetic error would not be expected to generate more evidence for co-divergence. For example, the grouping of kolmioviruses from placental mammals, marsupials, birds and reptiles exactly replicated the host phylogeny. It is likely that additional co-divergence events will be revealed by more sampling.

Critically, irrespective of which phylogenetic tree or data set is used, the most distinctive aspect of the *Kolmioviridae* phylogeny is that individual host groups contain multiple and highly distinct lineages. This evolutionary pattern is best documented by the four highly distinct lineages (likely representing different genera) found in crocodilians and reptiles, and the three distinct clades found in birds that cover the entire genetic diversity of the *Kolmioviridae*. Hence, rather than viewing kolmiovirus evolution as a single lineage characterized by multiple co-divergence events and/or host species jumps, our phylogenetic analysis indicates that multiple distinct kolmiovirus lineages have co-circulated in vertebrates over timescales of hundreds of millions of years. The presence of these co-circulating lineages raises additional questions that are beyond the scope of this study. For example, do these lineages differ in fundamental biology? Are they competition with each other? Additional sampling of kolmioviruses in nature will help answer these questions.

Where virus-host co-divergence does occur on the kolmiovirus phylogeny we are able to add a tentative evolutionary timescale, utilizing established dates of host divergence. Accordingly, the divergence between the mammalian and the reptile/bird lineages (e.g., node 1, Fig. 1) would date to approximately 310 million years ago, while that of fish from other vertebrates (node 4) approximately 450 million years ago. Hence, we may conclude that the vertebrate kolmioviruses are at least 450 million years old. However, it is clear that there is no overall molecular clock of kolmiovirus evolution, with substantial variation in branch lengths which will greatly distort estimates of divergence times. As a case point, the genetic diversity within the diversity of human HDV sequences is evidently far greater than if it only reflected the diversification and dispersal of anatomically modern humans over the last 200,000 years (Fig. 1).

Also of note in this context is that there is currently no evidence for the presence of kolmioviruses in non-human primates, in turn suggesting that human HDV originated via a cross-species transmission event from another mammalian host. However, it is unclear when, where, and from what animal reservoir this cross-species transmission event occurred, and there is a major temporal mismatch between the HBV and HDV phylogenies. In particular, given that ancient DNA data suggests that the known genotypes of human HBV are no older than approximately 15,000 years (24) with no evidence for co-divergence among primates (i.e., the HBVs found in African and Asian great apes fall within the diversity of human sequences), it seems likely that the association between HBV and HDV itself only occurred in the last 15,000 years or so, and that the 19-20 amino acid extension to the C terminal of the LDAg occurred at this time. As well as a functional compatibility, the high prevalence of HBV infection in humans, its long duration of infection within individual hosts, and its propensity to generate surface antigen particles (HBsAg), meant that it could provide a ubiquitous substrate for HDV replication and transmission. By leading to an increased number of transmission events, and hence replications per unit time, the association between HBV and HDV may also have increased the rate of evolutionary change in the latter, perhaps explaining why human HDV appears proportionally more divergent than other vertebrate kolmioviruses.

Overall, our study suggests that kolmioviruses have been in existence for the entire duration of vertebrate evolution, reflecting an evolutionary process that involves the co-circulation of multiple viral lineages for extended time periods. Over this extended timescale kolmiovirus evolution has been characterized by a complex mix of co-divergence and cross-species transmission, a macroevolutionary pattern that now appears to be true for many RNA viruses. Undoubtedly, a better understanding of kolmiovirus evolution, with increased phylogenetic resolution, will rest on the successful identification of kolmioviruses in a broader range of vertebrate taxa along with detailed experimental assessment of the functional relationships between kolmioviruses and diverse helper viruses.

## Materials and Methods

### Animal Ethics

American alligator (*Alligator mississippiensis*) sample collection was approved by the Clemson University Animal Use Protocol #2022-0454 and through research permits from the South Carolina Department of Natural Resources (Rainwater permit 2024), Georgia Scientific Collections Permit (No. 1001742080), Savannah National Wildlife Refuge (Permit No. 2025-16), Florida Fish and Wildlife Conservation Commission (No. SPGS-25-44), and Alabama Scientific Collection Permit License (No. 2024149244868680). Alligator sampling in North Carolina was performed with the North Carolina Wildlife Resources Commission requiring no specific permits. Red kangaroo (*Osphranter rufus*) samples were collected under the ethics approval 4b/10/24 provided by Taronga Conservation Society Australia’s Animal Ethics Committee and under the auspices of scientific license SL100104 issued by the New South Wales Department of Energy, the Environment, Climate Change and Water. Samples from the central bearded dragon (*Pogona vitticeps*) were obtained from a captive breeding colony derived from a wild population under ethics approval (AEC13401) from the University of Canberra.

### Sampling and sequencing of American alligator (Alligator mississippiensis) samples

Alligator samples were opportunistically collected between April 2024 to May 2025 from 22 sites and across five states in the USA: North Carolina, South Carolina, Georgia, Florida, and Alabama (Table S1). All samples were preserved in an ammonium sulphate and sodium citrate-based RNA preservation buffer (prepared in-house as an RNAlater alternative), kept on ice during transport and stored at −80°C until sample extraction. Tissue samples from hunter-harvested alligators were taken post-mortem and kept in 2mL RNAlater buffer, while blood samples were taken in the ratio 1mL blood to 1mL RNAlater, and sterile FLOQSwabs (Copan, California, USA) were used for oral and cloacal samples in 1mL RNAlater. All samples were extracted using the QIAShredder and QIAGEN RNeasy Plus Mini Kit (QIAGEN, Hilden, Germany) following the manufacturer’s protocol and kept frozen at −80°C until transport to the sequencing facility overnight on dry ice.

Quality control and total RNA sequencing were conducted by the Yale Center for Genome Analysis (YCGA; Yale University, Connecticut, USA). Libraries were prepared using the WatchMaker RNA library prep kit with Polaris rRNA depletion (Watchmaker Genomics, Colorado, USA). A total of 31 libraries were sequenced using the NovaSeqX Plus platform (150bp paired end; Illumina, San Diego, USA), of which three contained evidence for kolmiovirus infection and were hence studied herein. We note that the negative control, generated by running buffers through clean kit columns, yielded insufficient RNA for library preparation and was therefore excluded from sequencing.

### Sampling and sequencing of red kangaroo (Osphranter rufus) and central bearded dragon (Pogona vitticeps) samples

Lung tissue from a red kangaroo was collected in collaboration with the Australian Registry of Wildlife Health, Taronga Conservation Society Australia (Sydney, NSW) (Table S1). The collection of lung and liver tissue from a central bearded dragon was obtained from a captive breeding colony held at the University of Canberra, with the individuals derived from a wild population (Table S1). Tissues were snap frozen at −80°C.

Kangaroo lung, and bearded dragon liver and lung samples were homogenized using TissueRuptor (QIAGEN, Hilden, Germany) and extracted using QIAGEN RNeasy Plus Mini Kit following the manufacturer’s protocol, and kept frozen at −80°C until transport on dry ice to the Australian Genome Research Facility (AGRF, Melbourne, Australia). rRNA was depleted using the Illumina rRNA depletion kit and libraries were prepared using the Illumina Truseq Total RNA Library Preparation Protocol prior to sequencing on the Illumina NovaSeqX platform (150bp paired end; Illumina, California, USA).

### Novel kolmioviruses in the Sequence Read Archive (SRA)

The Serratus Explorer (10) was used to screen for novel kolmioviruses in the SRA. Accordingly, a total of 57 non-human SRA libraries were detected to contain kolmioviruses (see Results). Of these 57 SRA libraries, 46 came from vertebrates and were subsequently downloaded using command lines adapted from the BatchSRAMiner script (https://github.com/JonathonMifsud/BatchArtemisSRAMiner). Libraries known to contain kolmioviruses and with hosts known to be associated with kolmioviruses were included as positive controls to validate the virus discovery pipeline (Table S2).

### De novo assembly of sequenced libraries and SRA libraries

Raw sequence reads were trimmed using Trimmomatic (v.0.35; 25) in the configuration “LEADING:3 TRAILING:3 MINLEN:36”. Raw and trimmed libraries were evaluated using FastQC (v.1.9.; 26) to confirm the quality of the reads and the successful trimming of adapter sequences. The metagenomic taxonomic classifier Kraken2 (v.2.1.3.; 27), with a confidence threshold set at 0.2 using the “core_nt” database (as of October 2025) was used to identify any potential library contamination.

Contigs were assembled using MEGAHIT (v.1.2.9; 28) with the read abundance of these contigs estimated using RNA-Seq by Expectation Maximisation (RSEM-v1.3.1; Bowtie2-v2.5.4; Salmon-v1.10.3; Samtools-v1.22.1). Contigs were first screened against the clustered Reference Viral Database (RVDB v29; 29) using DIAMOND BLASTx (v.2.16.0.; 30) with the flags set to “-e 1E-5 - -ultra-sensitive”. Potential virus contigs were then screened against the BLAST Non-Redundant Protein DIAMOND database (as of August 2025) to eliminate false-positive contigs. Putative kolmiovirus contigs were then subjected to open reading frame (ORF) prediction using Geneious (v.2025.1.) and a manual web-based BLAST analysis of the delta antigen (DAg) for kolmiovirus confirmation.

Reference kolmiovirus sequences from previous studies (9-12, 31; Table S3), as well as those available on NCBI/GenBank, were curated to best represent the diversity and distribution of kolmioviruses among vertebrates. To assess the placement of vertebrate-associated kolmioviruses relative to those detected in invertebrates, the termite HDV-like virus (GenBank accession NC_076410) was included in some analyses (Fig. S1). Human kolmiovirus (deltavirus) sequences were curated to represent all recognized species/genotypes. As kolmioviruses possess circular genomes and given the need for multiple sequence alignment prior to phylogenetic analyses, all genomes were circularized and then linearized at the DAg start codon. Only sequences with complete DAg ORFs were included in the phylogenetic analysis. This resulted in a total of 54 sequences used to construct vertebrate-associated kolmiovirus phylogenetic trees. Full and partial nucleotide sequences ranged between 564 nucleotides (BR001660 Erythrura gouldiae-associated kolmiovirus) and 2415 nucleotides (Red Kangaroo kolmiovirus).

### Phylogenetic analysis

The kolmiovirus sequences newly generated here and those previously identified were aligned using three methods: (i) Clustal Omega (32), (ii) MUSCLE (33) and (iii) MAFFT (34). In some cases, sequence alignments were trimmed by removing ambiguously aligned sites using trimAl (35) with the -gappyout flag (see Results).

Phylogenetic trees of these alignments were then estimated using the maximum likelihood (ML) method in IQ-TREE2 (36) with the ModelFinder Plus (-m MFP) flag used to select the best-fit models of amino acid and nucleotide substitution (37). Different phylogenetic trees were estimated to assess the impact of different (i) data sets (full genome, DAg), (ii) sequence alignment methods (Clustal Omega, MUSCLE, MAFFT), (iii) trimAl trimmed and untrimmed outputs, and (iv) nucleotide and amino acid substitution models (best-fit model based on ModelFinder Plus Bayesian Information Criterion, Akaike Information criterion, free rate models, simpler user-defined substitution models) on topological structure, particularly the extent of virus-host co-divergence. After model testing, the following substitution models were utilized: (a) full genome nucleotide sequences were analyzed using the Three-Parameter Model 2 model with AC=AT, AG=CT, CG=GT and incorporating invariable sites and a FreeRate model of rate heterogeneity with three categories (i.e., the TPM2+I+R3 substitution model); (b) DAg nucleotide sequences were analyzed using the Transitional Model 2 with AC=AT and CG=GT, empirical base frequencies, and a gamma-distributed rate heterogeneity (TIM2+F+Γ4); and (c) DAg amino acid sequences were analyzed using the insect-derived Q matrix with empirical amino acid frequencies and gamma-distributed rate heterogeneity (Q.insect+F+Γ4) (36). In all phylogenies, nodal support was estimated using 1000 bootstrap replicates. Phylogenies were further annotated using FigTree (v1.4.4) and edited using Adobe Illustrator (2025).

The number of co-divergence events in each phylogeny was counted as the number of nodes from (i) virus clades delineating different groups of vertebrates when there was more than one virus in that clade (i.e., humans, placental mammals, marsupials, birds, amphibians, reptiles) summed with (ii) the number of nodes that then linked these vertebrate groups in a manner that matched the host phylogeny.

### Phylogeny based on delta antigen structural prediction

DAg protein structures were predicted from amino acid sequences using AlphaFold3 seeded at 1000 (38). High confidence models were selected based on the AlphaFold3 scoring index which utilizes a per-residue confidence score measured by the predicted Local Distance Difference Test (pLDDT) and overall structural confidence defined by pTM/ipTM. The mean pLDDT across all models was 69%. Models were aligned with FoldMason (39) using the command ‘--match-ratio 0.9 --filter-msa 1 --gap-open aa:25,nucl:25 --gap-extend aa:2,nucl:2 --report-paths 0 --report-mode 2’. The resultant FoldMason Multiple Structural Alignment (MSTA) maximizes the average Local Distance Difference Test (LDDT) score as a structure similarity metric and was reported with a final value of 0.472. The MSTA was then subject to trimming of columns containing ≥35% gaps using trimAl (35). ML phylogenetic analysis of the MSTA was conducted using IQ-TREE2 (36) with ModelFinder Plus (-m MFP) and 1000 normal bootstraps on the 3Di and amino acid MSTA. A 3Di and amino acid edge-linked partitioned tree was also estimated using the FoldMason outputs, where the 3Di sequence utilized a custom 3Di substitution matrix (40) and the amino acid sequence alignment employed the Q.insect+F+Γ4 substitution model as defined by ModelFinder. Structural visualization and analysis were performed using USCF ChimeraX (v1.11.1; 41). To confirm that low pLDDT regions corresponded to intrinsically disordered regions (IDR), protein disorder in the novel kolmioviruses was predicted using AIU-Pred (v2.2.1; 42).

## Data availability

The raw sequence reads generated in this study are available at the NCBI SRA under BioProject PRJNA1474605 accessions SRR38998513-SRR38998518. Virus consensus sequences have been deposited in the NCBI GenBank database under accession numbers XXX-YYY.

## Acknowledgments

This work was funded by a National Health & Medical Research Council (Australia) Investigator grant (GNT2017197) and an Australian Research Council Discovery Project grant (DP240101313) to ECH. The authors acknowledge the technical assistance provided by the Sydney Informatics Hub at the University of Sydney. This work was also supported in part by the\ U.S. Department of Energy (DE-EM0004391) in association with SREL and represents technical contribution number 7554 of the Clemson University Experimental Station.

Alligator sample collection was supported by state agency and private collaborators in North Carolina, South Carolina, Georgia, Alabama, and Florida. We appreciate the assistance of: Alicia Wassmer (North Carolina Wildlife Commission), residents of Camp Bryan (Lake Ellis-Simon, NC), Morgan Hart (South Carolina Department of Natural Resources), Jamie Dozier (Tom Yawkey Wildlife Center), Dr. Andrew Bridges and Beau Bauer (Nemours Wildlife Foundation), Tony Mills (Spring Island Trust), Brandon Goff (Ramsey Grove State Park), Chuck Hayes (Savannah National Wildlife Refuge), Kara Nitschke and Theron Menken (Georgia Department of Natural Resources), Joseph Colbert and Yank Moore (Jekyll Island Authority), Mike Gifford (“Gator Mike”), Richard Tharp (Alabama Department of Conservation of Natural Resources), Dr. Chris Murray (Southeastern Louisiana State University), Mike Haley, Matt Nichols (Florida Fish and Wildlife Commission Alligator Management).

## Author Contributions

L.L. and E.C.H. conceptualized the study. M.B., S.L.W., and K.R. performed the animal sampling. L.L., K.V.B., J.M., and E.M. analyzed the data. T.R.R., J.T.A., B.B.P. and E.C.H. provided supervision. L.L. and E.C.H. wrote the original draft. K.V.B., M.B., B.B.P., S.L.W., K.R. provided manuscript revisions and additions. All authors reviewed and approved the final manuscript.

## Competing Interest Statement

The authors declare no completing interests.

## Notes

### Competing Interest Statement

The authors have declared no competing interest.

### Summary of Updates

1. Change of title from "Kolmioviridae" to "Kolmiovirus" 2. Minor text changes 3. Removal of Table S4 (originally included by mistake)

