## Supplementary material for "Co-Circulation of Multiple Kolmiovirus Lineages Through Vertebrate Evolution": Figures S1-S3

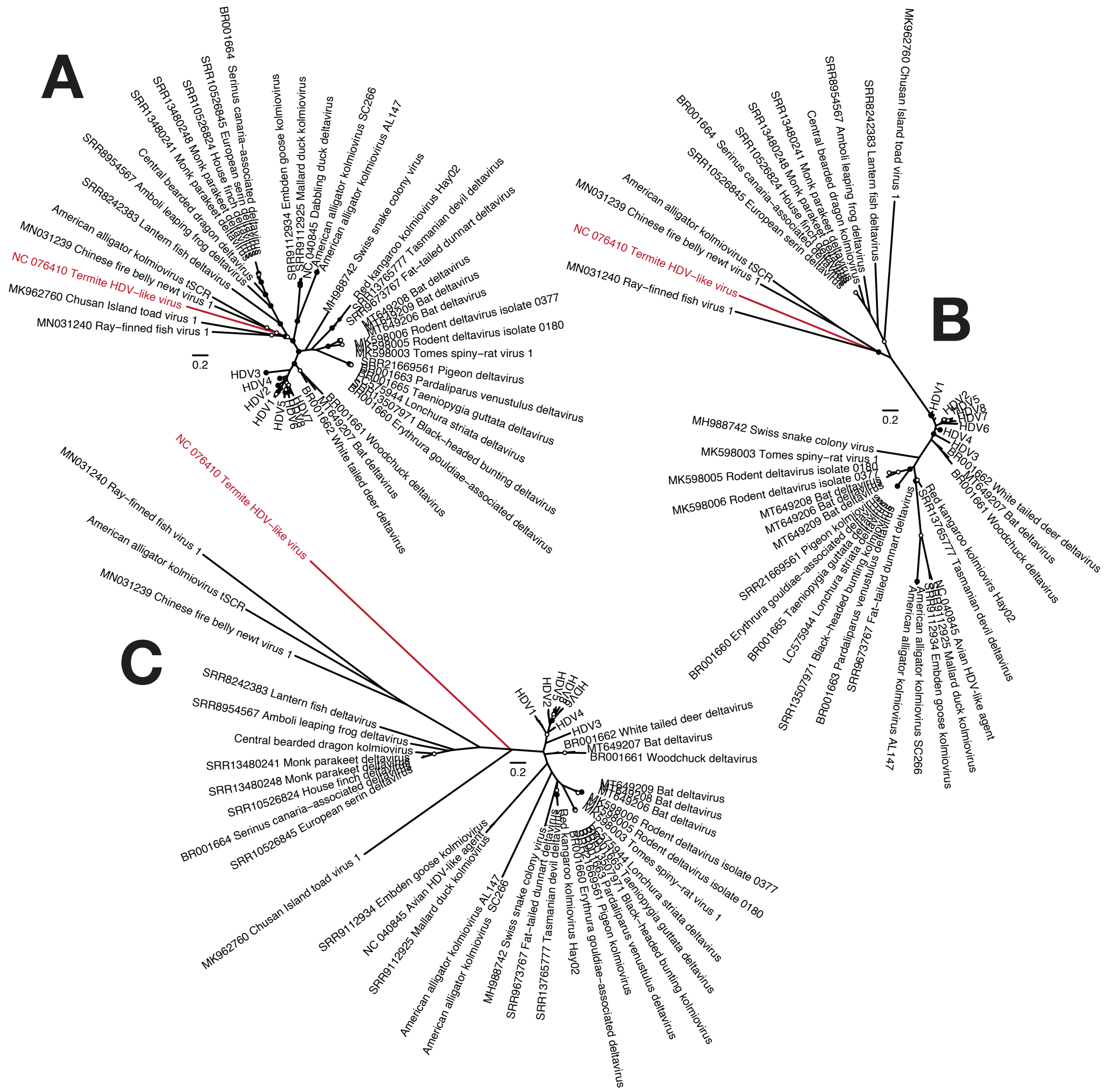


**Supplementary Figure 1. Maximum likelihood phylogenetic trees of vertebrate host-associated *Kolmioviridae* with a termite (accession NC_076410) outgroup.** All ML trees were estimated using Clustal Omega, trimAl, ModelFinder (see Table 1 for details), with 1000 bootstrap replicates; nodes ≥95% bootstrap support are annotated with black circles (●) and those with 70-94% bootstrap support with white circles (○). (**A**) ML phylogeny based on the nucleotide sequence of the complete viral genome. (**B**) ML tree based on LDAg nucleotide sequences. (**C**) ML tree based on LDAg amino acid sequences.

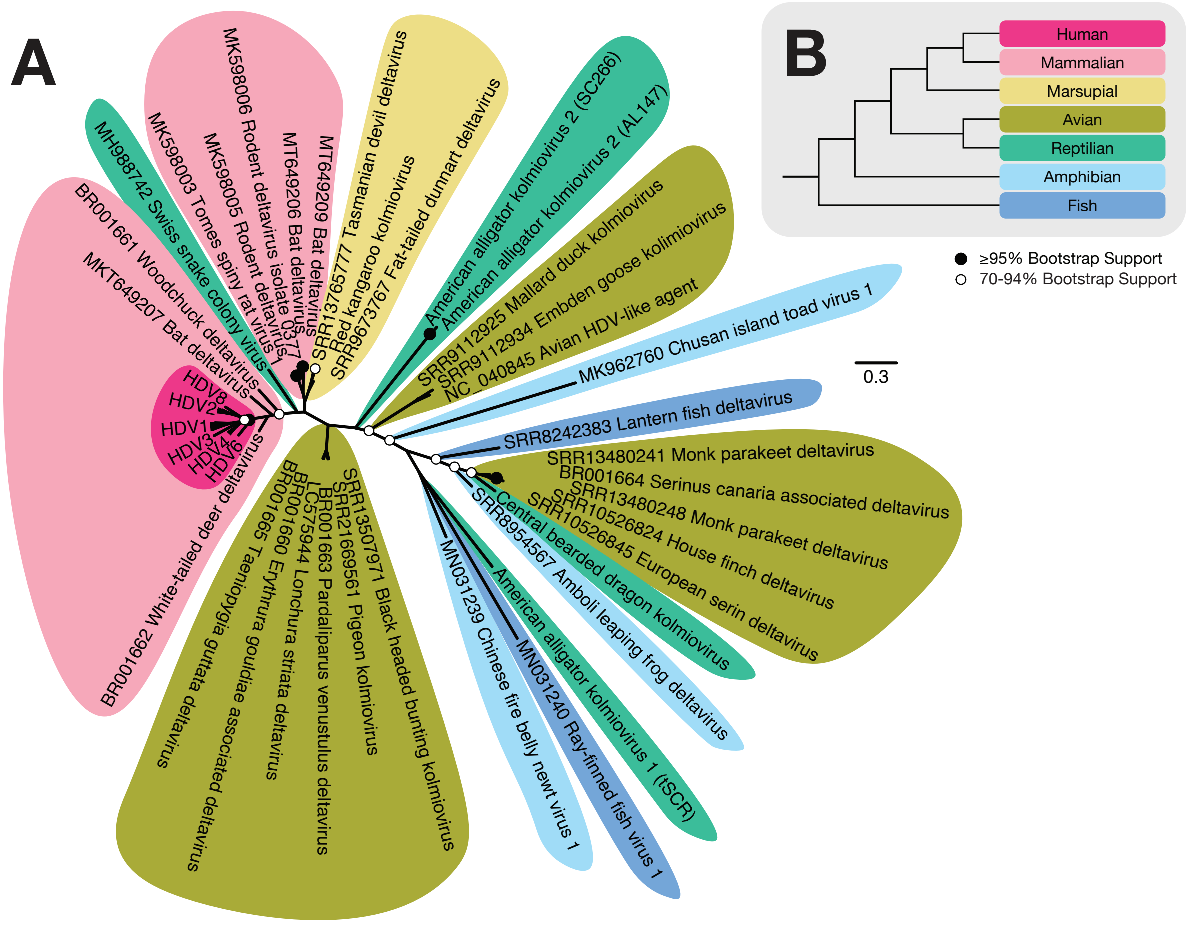


**Supplementary Figure 2. AlphaFold3 predicted structural phylogeny.** (**A**) Unrooted ML phylogenetic tree of vertebrate host-associated *Kolmioviridae* estimated using the FoldMason Multiple Structural Alignment (MSTA) from which a 3Di+AA partitioned tree was inferred. A custom 3Di substitution matrix was utilized for 3Di and ModelFinder (Q.insect+F+$\Gamma_4$) for amino acid sequences (see Table 1 for details). Nodal support estimated using 1000 bootstrap replicates; nodes ≥95% bootstrap support are annotated with black circles (●) while those with 70-94% bootstrap support are shown with white circles (○). (**B**) Phylogeny of major vertebrate host taxa colour key: human (hot pink), placental mammals (light pink), marsupials (yellow), birds (army green), reptilians (green), amphibians (light blue), fish (dark blue).

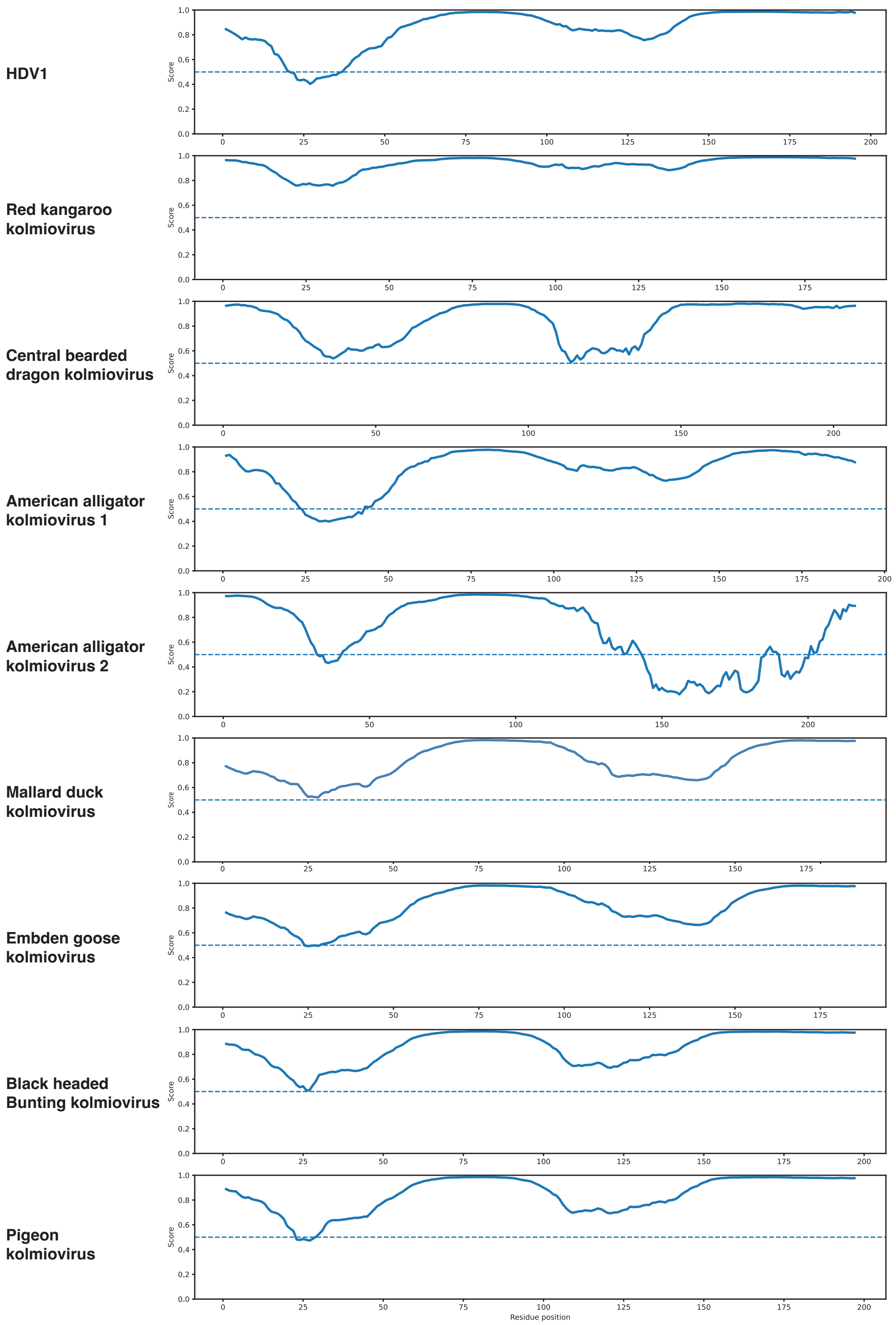


**Supplementary Figure 3. Predicted intrinsic disorder in novel kolmioviruses showing high disorder in regions of low pLDDT.** Disorder prediction and scoring using AIU-Pred and delta antigen amino acid sequences. Regions of reduced disorder correlate with greater prediction confidence (pLDDT >70%) and correspond to conserved structural domains, including coiled-coil and helix-loop-helix.
